# Increasing Synaptic GluN2B levels within the Basal and Lateral Amygdala Enables the Modification of Strong Reconsolidation Resistant Fear Memories

**DOI:** 10.1101/537142

**Authors:** Christopher A. de Solis, Cuauhtémoc U. Gonzalez, Mario A. Galdamez, John M. Perish, Samuel W. Woodard, Carlos E. Salinas, Joel N. Miller, Hajira Elahi, Oziel H. Pineda, Shayrin Oad, Serafin Gatica de las Fuentes, Malissa S. Owen, Alfredo Sandoval, Roopashri Holehonnur, Jonathan E. Ploski

## Abstract

Reconsolidation disruption has been proposed as a method to attenuate pathological memories in disorders such as PTSD. However, studies from our group and others indicate that strong memories are resistant to becoming destabilized following reactivation, rendering them impervious to agents that disrupt the re-stabilization phase of reconsolidation. Thus, therapies designed to attenuate maladaptive memories by disrupting reconsolidation updating have not been adequately developed. We previously determined that animals possessing strong auditory fear memories, compared to animals with weaker fear memories, are associated with an enduring increase in the synaptic GluN2A/GluN2B ratio in neurons of the mouse basal and lateral amygdala (BLA). In this study, we determined whether increasing GluN2B levels within BLA excitatory neuronal synapses is sufficient to enable modification of strong fear memories via reconsolidation. To accomplish this, we utilized a combinatorial genetic strategy to express GluN2B or GluN2B(E1479Q) in excitatory neurons of the mouse BLA before or after fear memory consolidation. GluN2B(E1479Q) contains a point mutation that increases synaptic expression of the subunit by interfering with phosphorylation-driven endocytosis. At the time of memory retrieval, increasing synaptic GluN2B levels by expression of GluN2B(E1479Q), but *not* GluN2B(WT), enhanced the induction of reconsolidation rendering the strong fear memory modifiable. GluN2B(WT) or GluN2B(E1479Q) expression did not influence fear memory maintenance or extinction. Fear memory consolidation, however, was enhanced when GluN2B(E1479Q) was expressed in the BLA at the time of training. These findings indicate that enhancing GluN2B synaptic trafficking may provide a novel therapeutic strategy to enhance modification of pathological memories.

## Introduction

Many previous findings indicate that memory retrieval (i.e., reactivation) of weak fear memories induces a period of vulnerability during which they are susceptible to modification or erasure. The prevailing view is that if these memories are not re-stabilized via the protein synthesis-dependent mechanism of reconsolidation, the memories are lost (1-4). This transient labilization is thought to allow new information to be effectively integrated into the existing memory trace (5, 6). These alterations may weaken or strengthen (i.e., modify) the original memory (7). Pharmacologically blocking the reconsolidation process is a potentially powerful treatment for attenuating memories associated with pathologies such as PTSD and drug addiction (8-11). Recent clinical evidence, however, indicates that the effectiveness of this approach may be limited because some memories are particularly resistant to retrieval-dependent memory destabilization (12). In these cases, memory retrieval does not initiate the reconsolidation process, rendering pharmacotherapies designed to disrupt reconsolidation ineffective (13,14). This indicates that reconsolidation consists of two distinct phases. The first phase, memory “destabilization,” is the initiation or induction of reconsolidation and represents the cellular and molecular events during memory retrieval that produce lability, or instability, in the memory trace. The second phase of reconsolidation is the protein synthesis-dependent re-stabilization, or re-storage phase, which can be targeted by blockers of reconsolidation (i.e., anisomycin, propranolol, mifepristone, etc.). It is now known that a number of variables, including the age of the memory, the strength of the memory, and the conditions under which memory retrieval occur, present important factors that may contribute to the effectiveness of memory retrieval to initiate memory destabilization and induce reconsolidation updating. Typically, strong or remote memories have been shown to be difficult to disrupt via reconsolidation blockade. This appears to be due to the resistance of these memories to undergo retrieval-dependent memory destabilization (13-16). Therefore, targeting the reconsolidation process will likely never be an effective treatment to attenuate pathological memories unless methods are devised to enhance the induction of their reconsolidation.

A number of studies currently indicate that there are at least three major cellular events that are critical for retrieval-induced destabilization of auditory fear memories: *N*-methyl D-aspartate receptor (NMDAR) activation (17, 18), ubiquitin proteasome-dependent protein degradation (19), and alterations in α-amino-3-hydroxy-5-methyl-4-isoxazolepropionic acid receptor (AMPAR) subunit trafficking/endocytosis (20). NMDAR activation is critical to initiate ubiquitin proteasome-dependent protein degradation and alterations in AMPAR subunit trafficking/endocytosis (19). NMDARs are heteromeric glutamate receptors composed of two obligatory GluN1 subunits and two GluN2 subunits (21-23). In the hippocampus and amygdala, the GluN2A and GluN2B subtypes are the predominantly expressed GluN2 subtypes (24, 25). NMDAR subunit composition alters the properties of the NMDAR due to inherent differences between the GluN2A and GluN2B subunits. GluN1:GluN2A receptors have a higher open probability, faster decay time, and lower sensitivity to glutamate and glycine than GluN1:GluN2B containing receptors. GluN2A and GluN2B also differ considerably in their ability to bind to postsynaptic scaffolding proteins that regulate the induction of plasticity (26, 27).

In an effort to understand the molecular basis of the differential requirements for retrieval-dependent memory destabilization between weak and strong fear memories, our laboratory recently determined that auditory Pavlovian fear memories created with 10 tone-shock pairings (10 TSP) are resistant to retrieval-dependent memory destabilization, and are associated with an increase in the synaptic GluN2A/GluN2B ratio in neurons of the BLA when compared to weaker fear memories created via 1 or 3 tone-shock pairings (1-3 TSP) (16). This switch could explain why strong fear memories are resistant to retrieval-dependent memory destabilization and modification via reconsolidation. Prior to this study, it was unknown whether increasing synaptic GluN2B levels within the BLA is sufficient to render strong fear memories modifiable via reconsolidation. Our findings indicate that manipulations that lead to an increase in synaptic GluN2B may be viable strategies to render pathological memories modifiable via reconsolidation updating.

## Methods

### Subjects

Two to three-month-old male and female C57BL/6 αCaMKII-tTA mice (Jackson Laboratories; cat # 003010 B6) (28) were used for experiments described in this study. Male and female mice were used in all experiments, and were equally distributed across the experimental groups. Animals were individually housed in polycarbonate cages on a 12-hour light/dark cycle. Food and water were provided ad libitum. Animal use procedures were conducted according to the National Institutes of Health Guide for the Care and Use of Laboratory Animals and were approved by the University of Texas at Dallas Animal Care and Use Committee.

### Lentiviral plasmids

Viral plasmids were created using standard recombinant cloning techniques. The development and testing of lentiviral plasmids encoding TRE3G-Flag-GluN2B and TRE3G-GFP are described in a previous study from our laboratory (29). The lentiviral plasmid encoding TRE3G-Flag-GluN2B(E1479Q) was created by excising a ∼1200 bp SbfI-SacII DNA fragment from the lentiviral plasmid encoding TRE3G-Flag-GluN2B and exchanging it with a similar DNA fragment (gBlock; Integrated DNA Technologies), which contained a codon mutation resulting in the E1479Q change. The single Flag tag in the lentiviral plasmids encoding TRE3G-Flag-GluN2B and TRE3G-Flag-GluN2B(E1479Q) was converted to a 4X Myc tag by excising a ∼240 bp BamHI-XmaI DNA fragment out of these plasmids and exchanging it for an ∼360 bp DNA fragment which replaced DNA coding for the Flag tag for DNA coding for a 4X Myc tag (gBlock; Integrated DNA Technologies). All plasmids were verified by DNA sequencing.

### Viral production, purification, and titering

Large-scale viral production, purification, and titering were performed as described previously (29).

### Fear Conditioning

Animals were fear conditioned and tested using a standard auditory fear conditioning system equipped with video monitoring (Coulbourn Instruments), as previously described (16). Habituation: Two days post-viral infusion, mice were handled for 2-3 min for two days. Mice were habituated to both the training context and retrieval context for 10 min per day for four days. For reconsolidation experiments, two days before training, mice were habituated to infuser placement. Training: Weak fear conditioning (3 TSP) consisted of exposure to 3 tones (30 sec, 5 kHz, 75 dB), each co-terminating with a 2 sec, 0.75 mA foot shock, with an inter-trial interval (ITI) of 110 sec. For strong fear conditioning (10 TSP), animals received 10 similar TSPs; however, the ITI of the 10 TSP was pseudorandom (3-7 min average ITI over 45 min). Forty-eight hours after fear conditioning, animals were exposed to a single 30 sec tone in an alternate context (i.e., a modified chamber and the absence of light, with distinct olfactory and tactile cues). Post-reactivation LTM (PR-LTM) was examined 24 hr after reactivation and consisted of 5 tones in the same context as reactivation. For consolidation experiments, short-term memory (STM) was examined 3 hours after training by exposing the mice to 3 tones (2 min ITI; 30 sec, 5 kHz, 75 dB) in an altered context. The long-term memory (LTM) test consisted of five 30 sec tones 24 hr following training in the same altered context as STM. Videos were saved to a hard drive and freezing to the tone was scored by individuals blinded to experimental conditions.

### Viral infusion and Cannula Implantation

Viral infusions targeted the BLA [AP −1.6, ML ±3.3, -DV ±4.97] of αCaMKII-tTA mice. Mice were anesthetized with ketamine (100 mg/kg) and xylazine (10 mg/kg). Infusion cannula (C315G; PlasticsOne) were inserted into tubing (I.D. 0.0150 in, O.D. 0.043 in, wall thickness 0.0140 in; A-M Systems, Inc.) which was fit securely to a syringe [Hamilton Company; 2 μL, 23-gauge (88500)]. Prior to surgery, the tubes were backfilled with sterile 1X PBS, followed by sesame oil, where only 1X PBS was present in the ∼5 cm region closest to the infusers. The virus was drawn up into the infusion cannula, infusers were lowered to the BLA of mice, and virus was infused (1 μL/side at a titer of 1E+9 GC/mL) at a rate of 0.07 μL/min for 15 min using an infusion pump (New Era Pump Systems Inc.; NE-300). For reconsolidation experiments, guide cannula (PlasticsOne; infuser projection, 1 mm; dummy cannula projection, 0.5mm) targeting the mouse BLA were implanted at the time of viral infusion and secured with dental acrylic (Ketac; Henry Schein Animal Health). To suppress the expression of the lentiviral transgene, animals were placed on a diet of 200 mg/kg Dox feed (BioServ) 2 days before surgery. All BLA lentiviral infusions contained a small amount of adeno-associated virus (AAV; 1E+8 viral particles) designed to express GFP from a cytomegalovirus (CMV) promoter to aid in viral placement analysis. Successful viral infusion and cannula placement were assessed by post-experimental perfusion of the animals with 10% phosphate-buffered formalin (110 mM, NaH2PO4, 150 mM NaCl) and observing GFP expression across the BLA (Bregma 2.3 to −0.58) in perfused sections. Sections were then stained with a 0.5% cresyl violet solution to aid in the identification of the loci of drug infusions. Only animals with bilateral BLA virus and cannula placement were used in the analyses.

### Drug Administration/Infusion

Following reactivation of the fear memory, mice were immediately removed from the behavior chambers and bilaterally infused with either anisomycin (125 μg/μL) or vehicle (0.9% sterile saline) to the BLA. Anisomycin or vehicle was infused at a volume of 0.3 μL/side at a rate of 0.15 μL/min. Anisomycin was prepared as we have previously (16).

### Post-synaptic Density Isolation

This procedure was performed as previously described (16). Mouse BLA transduced with lentivirus were dissected. To accomplish this, three 200 μm coronal sections containing the BLA were taken using a cryostat. BLA tissue was subsequently collected using a 0.5 mm punch tool. This tissue was Dounce homogenized in a Tris-HCl buffer (30 mM Tris-HCl, 4 mM EDTA, 1 mM EGTA). A protease inhibitor cocktail (Roche; cOmplete ULTRA Tablets) was used per manufacturer’s instructions in all PSD preparation buffers. The homogenate was centrifuged for 10 min at 800 G. The supernatant was collected and centrifuged for 1 hr at 100,000 G. The resulting pellet was resuspended in Tris-HCl buffer containing 0.5% Triton X-100 and incubated on ice for 20 min. Samples were then layered over 100 μL 1 M sucrose and centrifuged for 1 hr at 100,000 G. The resulting pellet was resuspended in Tris-HCl buffer containing 1% SDS. The concentrations of the samples were obtained by comparison to BCA standards (Pierce BCA Protein Assay Kit; 23227).

### Immunocytochemistry (ICC), Immunohistochemistry (IHC), and Western Blot

Immunostaining for Flag in 293FT cells and the mouse brain was carried out as previously published (29). Samples stained using ICC and IHC were imaged at 200X magnification (Olympus BX51 microscope, Olympus DP71 Digital Camera, and DP manager software). Western blot was performed on post-synaptic densities (PSDs) isolated from the mouse BLA. Conditions for GluN2B and PSD-95 were as described previously (16). Myc was detected using an anti-Myc monoclonal antibody (ThermoFisher; MA1-980). PVDF membranes containing the samples were blocked for 1 hr at RT using 5% BSA in TTBS. Samples were incubated overnight at 4° C with the anti-Myc antibody (1:1000) in 5% BSA in TTBS. Samples were washed 3 times in TTBS and incubated with anti-mouse-HRP (Cell Signaling, 7076S; 1:10,000) in 5% BSA for 1 hr at RT.

### Statistics

Analysis of western blot data was performed using a One-way ANOVA and a One-tailed *t*-test. One-way ANOVA was used to measure differences across groups during reactivation. One-way ANOVA with repeated measures was used to measure differences across groups during extinction and consolidation experiments. Two-way ANOVA with repeated measures was utilized to detect virus X treatment (anisomycin or vehicle) interaction effects in tests of PR-LTM. Fisher’s LSD was used for *post hoc* analysis of data analyzed with ANOVA.

## Results

### Lentiviruses designed to express GluN2B, GluN2B(E1479Q), or GFP do so *in vitro* and *in vivo*, and this expression can be suppressed using doxycycline

The goal of these experiments was to determine if increasing the level of GluN2B at BLA excitatory synapses at the time of reactivation of a strong fear memory is sufficient to enhance the retrieval-dependent initiation of reconsolidation. To accomplish this, it was necessary to genetically engineer recombinant viruses capable delivering GluN2B transgenes and to restrict expression of these transgenes to BLA excitatory neurons. We previously determined that GluN2B transgenes are difficult to express. Therefore, we developed an optimal method to increase GluN2B levels in αCaMKII positive BLA excitatory neurons by the infusion of lenti viruses into the BLA of αCaMKII-tTA mice. These viruses are designed to express GluN2B from a TRE3G promoter which restricts expression to αCaMKII-positive BLA excitatory neurons (when used within αCaMKII-tTA mice). These vectors allow sufficient expression of the transgene, and allow the transgene expression to be suppressed by feeding the animals doxycycline (Dox) containing chow (29).

Since it was previously reported that synaptic GluN2B levels may not be significantly influenced by mere overexpression of GluN2B mRNA/protein (30, 31), we generated a GluN2B viral transgene that contains a point mutation resulting in an E to Q transition within its C-terminal tail (E1479Q). This point mutation leads to inhibition of phosphorylation driven-endocytosis of GluN2B, leading to higher surface levels of the receptor subunit (32). We also generated a lentivirus to deliver green fluorescent protein (GFP) controlled from a TRE3G promoter as a control (**Figure 1a**).

**Figure 1.**
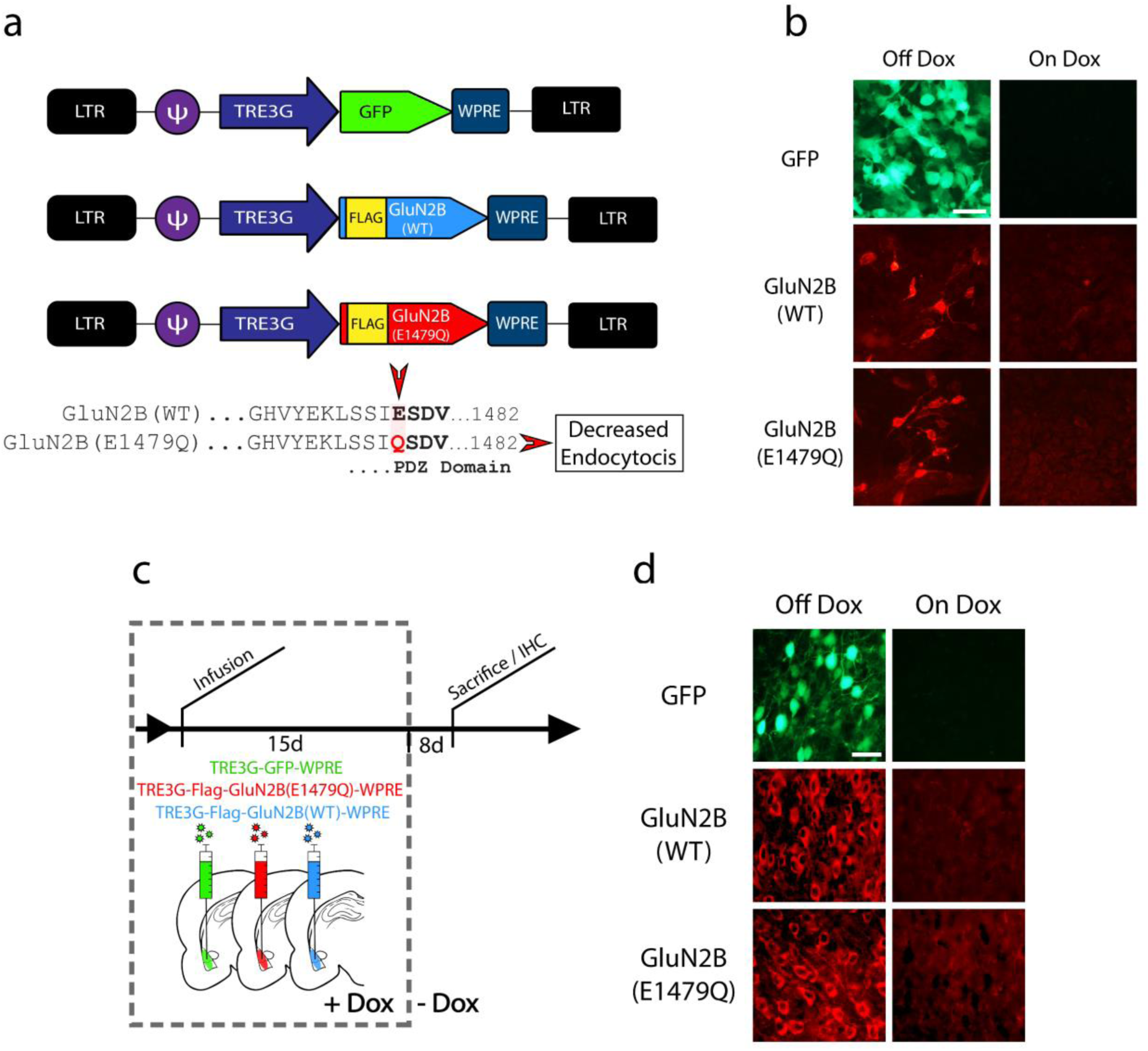
Lentiviruses designed to express GluN2B, GluN2B(E1479Q), or GFP do so *in vitro* and *in vivo*, and this expression can be suppressed using doxycycline. (a) Lentiviral genomes from 5’-3’ LTR (above). Location of mutation in the PDZ domain in GluN2B(E1479Q) compared to the GluN2B wild-type sequence (below). (b) Regulation of GFP, GluN2B(WT) and GluN2B(E1479Q) lentiviral transgenes by Dox *in vitro* (HEK cells) detected by imaging of GFP or immunocytochemistry for a Flag-tag included in each GluN2B transgene off Dox (left) and on Dox (right). (c) Timeline for testing Dox regulation of lentiviral transgenes *in vivo* in αCaMKII-tTA mice. (d) Detection of lentiviral transgene expression *in vivo* off Dox (left) and on Dox. All scale bars are set to 100 µm.

To determine if the expression of the lentiviral transgene is inhibited by the presence of Dox, 293FT cells expressing tetracycline-controlled transactivator (**tTA**) were transduced with these viruses in the presence and absence of Dox. Forty-eight hours post-transduction, an ICC was performed to examine Flag-tagged GluN2B(WT) and GluN2B(E1479Q) expression. GFP native fluorescence was also examined from the control construct. These viruses were capable of expressing their transgenes in the absence of Dox, and expression was suppressed in the presence of Dox (**Figure 1b**).

To determine if the pattern of expression was the same *in vivo*, we infused these lentiviruses into the BLA of αCaMKII-tTA mice while the animals were on Dox. We then removed half the animals from Dox and the animals were euthanized 8 days later. BLA-containing coronal brain sections were processed for detection of native GFP expression and IHC detection of the Flag-tagged GluN2B transgenes (**Figure 1c**). The viruses were capable of expressing their transgenes in the absence of Dox, and expression was suppressed in the presence of Dox *in vivo* (**Figure 1d**).

Next, we sought to assess if the delivered GluN2B(WT) and GluN2B(E1479Q) transgenes lead to incorporation of these proteins into the post-synaptic density (PSD) *in vivo*. For these experiments, we utilized 4X Myc-tagged GluN2B transgenes because we found that the Flag-tagged GluN2B transgenes were difficult to detect using anti-Flag western blotting in our pilot experiments (**Figure 2a**). We infused these viruses into the BLA of αCaMKII-tTA mice and harvested the tissue 15 days later (**Figure 2b**). The BLA was micro-dissected and the PSD proteins were isolated. Western blotting for Myc, GluN2B, and PSD-95 was subsequently performed. Samples were analyzed relative to PSD-95 expression. A One-way ANOVA revealed that GluN2B levels significantly differed between the GluN2B(WT), GluN2B(E1479Q), and Naïve samples, [*F*(2,25) = 3.55, *p* = 0.0461], [GluN2B(E1479Q) *n* = 9, GluN2B(WT) *n* = 9, Naïve *n* = 10]. *Post hoc* analysis determined that synaptic GluN2B levels were significantly higher in the GluN2B(E1479Q) samples compared to naïve animals (*p* = 0.0239). However, GluN2B levels were not significantly higher in the GluN2B(WT) samples compared to the naïve controls (*p* = 0.558), (**Figure 2c**). Western blotting for the Myc-tag indicated that both GluN2B(WT) and GluN2B(E1479Q) successfully incorporate into synaptic NMDARs at the PSD, but GluN2B(E1479Q) exhibited significantly higher synaptic levels compared to GluN2B(WT), [*t*(17) = 2.06, *p* = 0.027] (**Figure 2d**).

**Figure 2.**
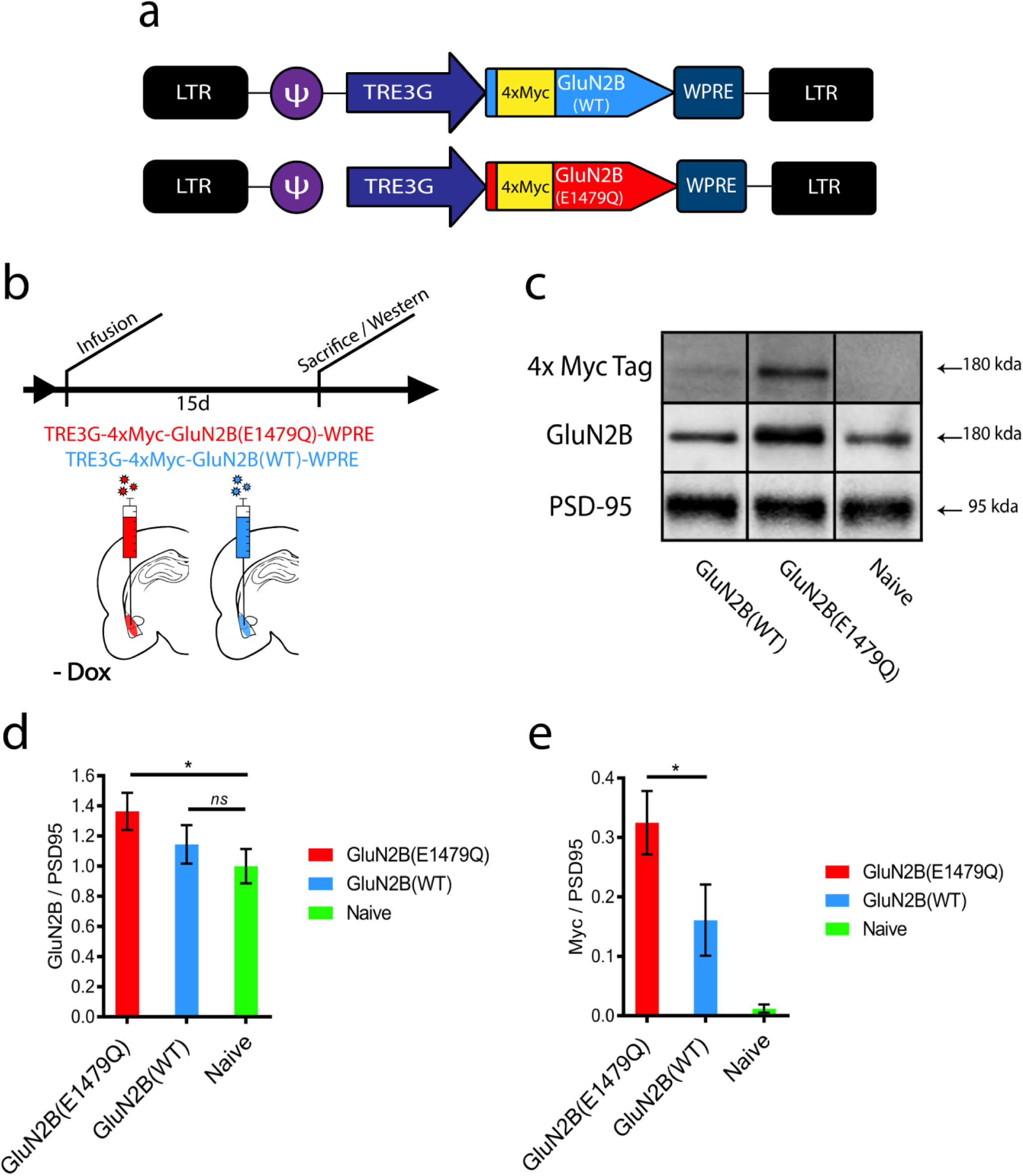
Overexpression of GluN2B via GluN2B(E1479Q), but not GluN2B(WT), increases levels of the subunit at the synapse. (a) Maps of lentiviral genomes designed to express GluN2B(WT) or GluN2B(E1479Q) with a 4X Myc-tag. (b) Experimental timeline for examining synaptic GluN2B levels within the BLA of αCaMKII-tTA mice following viral transduction with lentiviruses designed to express GluN2B(WT) or GluN2B(E1479Q). (c) Western blot of post-synaptic density proteins from amygdala neurons expressing GluN2B(WT), GluN2B(E1479Q), or naïve samples for Myc, GluN2B, and PSD95. (d) GluN2B levels quantified and compared to naïve samples reveals an increase in samples expressing GluN2B(E1479Q) (*p* < 0.04), but not in those overexpressing GluN2B(WT). (e) Myc signal quantified across groups reveals a significant increase of GuN2B incorporation into the post-synaptic density in samples that received GluN2B(E1479Q) compared to GluN2B(WT) (*p* < 0.03). * = *p* < 0.05, *ns* = not significant. Error bars = Standard Error of the Mean (SEM).

Collectively, these data are consistent with previous findings indicating that acute overexpression of GluN2B(WT) does not lead to significant differences in synaptic GluN2B levels. Synaptic GluN2B levels are regulated at the level of synaptic trafficking more so than on transcriptional/translational levels (30, 31). Thus, we can circumvent this limitation by utilizing GluN2B(E1479Q) to increase synaptic GluN2B levels.

### Increasing synaptic levels of GluN2B in BLA αCaMKII positive neurons using GluN2B(E1479Q) enables the reconsolidation of strong auditory fear memories and is retrieval-dependent and resistant to spontaneous recovery

In our next experiment, we infused the GluN2B(WT), GluN2B(E1479Q), or GFP viruses into the BLA of αCaMKII-tTA mice, and maintained the animals on a diet containing Dox to suppress the expression of the transgenes. The animals were fear conditioned with 10 TSP to produce strong fear memories. Two days later, the animals were switched to a diet that did not contain Dox to allow expression of the transgenes outside of the memory consolidation window. Eight days later, the animals underwent a single tone retrieval trial followed by immediate intra-BLA administration of vehicle or anisomycin. Twenty-four hours later, they were tested for PR-LTM (**Figure 3a**). GluN2B(WT) and GluN2B(E1479Q) were tested in separate experiments and compared to a GFP control. For the GluN2B(WT) experiment, there was no significant difference in freezing levels to the tone during reactivation across the groups, [*F*(1,24) = 0.065, *p* = 0.8002], [GFP/Veh *n* = 6, GFP/Aniso *n* = 7, GluN2B(WT)/Veh *n* = 7, GluN2B(WT)/Aniso *n* = 9], (**Figure 3b**). A Two-way ANOVA with repeated measures did not reveal a significant difference between the groups during PR-LTM, indicating that overexpression of GluN2B(WT) was not sufficient to enhance retrieval-dependent destabilization for a strong fear memory, [*F*(1,24) = 0.952, *p* = 0.3999]. For the GluN2B(E1479Q) experiment, there was not a significant difference in freezing levels to the tone during reactivation across the groups, [*F*(1,25) = 0.004, *p* = 0.9522], [GFP/Veh *n* = 7, GFP/Aniso *n* = 7, GluN2B(E1479Q)/Veh *n* = 7, GluN2B(E1479Q)/Aniso *n* = 7], (**Figure 3c**). However, a Two-way repeated measures ANOVA, [*F*(1,25) = 7.845, *p* = 0.0097], revealed a virus X drug interaction effect during PR-LTM. *Post hoc* analysis revealed that the GluN2B(E1479Q)/Aniso group froze significantly less than the other groups, (*p* < 0.05). This indicates that expression of GluN2B(E1479Q) was sufficient to enhance retrieval-dependent destabilization for a strong fear memory and render it vulnerable to reconsolidation blockade via post-retrieval intra-BLA anisomycin administration.

**Figure 3.**
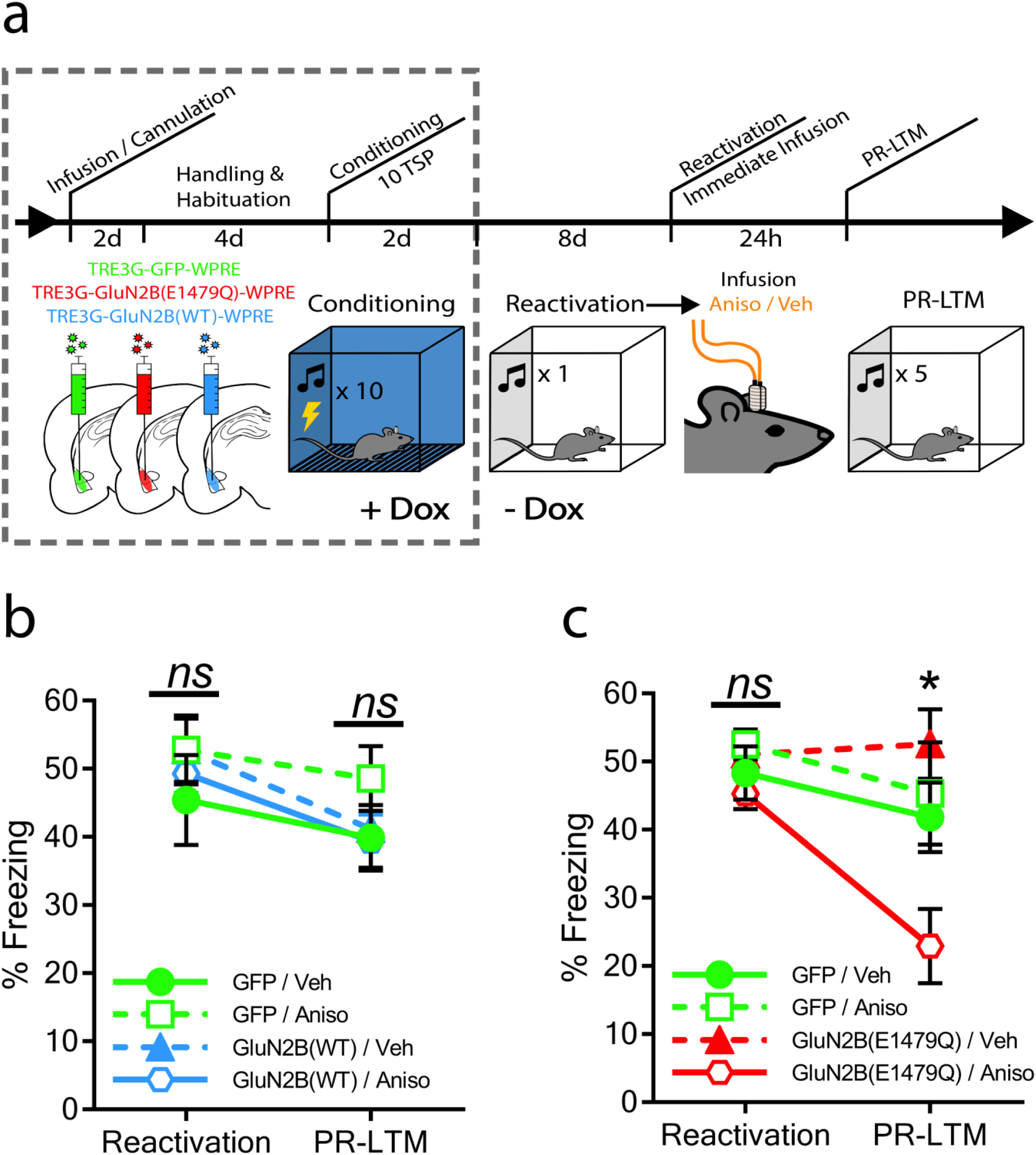
Increasing synaptic levels of GluN2B in αCaMKII positive BLA neurons using GluN2B(E1479Q) *but not* GluN2B(WT), enables the reconsolidation of strong auditory fear memories. (a) Experimental timeline of reconsolidation experiment. (b) Expression of GluN2B(WT) in BLA αCaMKII positive neurons was not sufficient to enhance retrieval-dependent destabilization for a strong fear memory. (c) Increasing synaptic levels of GluN2B in the BLA with GluN2B(E1479Q) at the time of reactivation of a strong fear memory did enhance the induction of reconsolidation to occur. Animals that received the mutant virus and were treated with anisomycin exhibited significantly lower freezing to the tone during a PR-LTM test (*p* < 0.001). * = *p* < 0.001, *ns* = not significant. Error bars = SEM.

As a follow-up to the above findings, a similar experiment was performed to assess if the apparent influence of GluN2B(E1479Q) on memory destabilization depends on retrieval of the fear memory. In this experiment, all animals were infused with GluN2B(E1479Q) virus in the BLA and maintained on a diet containing Dox. The animals were fear conditioned with 10 TSP 6 days after surgery. Two days later, the animals were switched to a diet that did not contain Dox. Eight days later, half the animals underwent a single tone retrieval trial and the other half did not. Animals in the retrieval and no-retrieval groups were then administered vehicle or anisomycin to the BLA. Twenty-four hours later, the animals were subjected to a PR-LTM test (**Figure 4a**). The animals that underwent reactivation did not exhibit a significant difference in freezing to the single tone, [*F*(1,15) = 0.809, *p* = 0.3826], (**Figure 4b**). A Two-way repeated measures ANOVA revealed a treatment X drug interaction effect between the groups during PR-LTM, [*F*(1,26) = 7.607, *p* = 0.0105]. *Post hoc* analysis revealed that mice that received both reactivation and anisomycin froze significantly less compared to the other groups (*p* < 0.05), [Reactivation/Veh *n* = 8, Reactivation/Aniso *n* = 9, No Reactivation/Veh *n* = 7, No Reactivation/Aniso *n* = 6]. In addition, we tested spontaneous recovery of fear 7 days following the PR-LTM test and were able to detect a statistically significant treatment X drug interaction effect in freezing to the tone, [*F*(1,26) = 8.227, *p* = 0.0081]. *Post hoc* analysis revealed that animals that received both reactivation and anisomycin froze significantly less compared to the other groups, (*p* < 0.05), indicating a lack of spontaneous recovery of freezing behavior. Collectively, these data indicate that GluN2B(E1479Q) mediated memory destabilization in a retrieval-dependent manner, and the reduction in freezing response due to the combination of GluN2B(E1479Q), reactivation, and anisomycin was resistant to spontaneous recovery when examined 8 days following reactivation.

**Figure 4.**
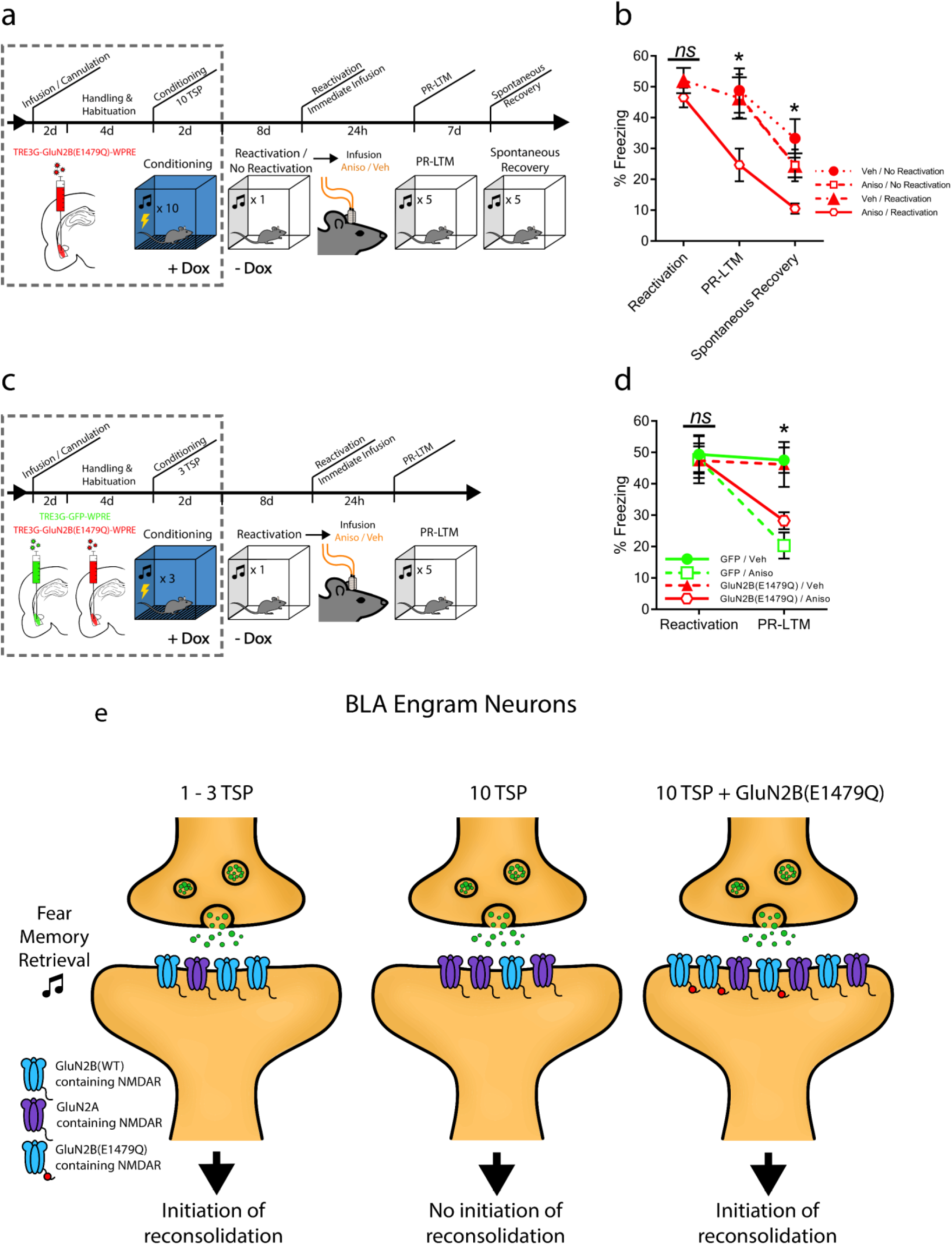
Increasing synaptic levels of GluN2B in αCaMKII positive BLA neurons using GluN2B(E1479Q) enables the reconsolidation of strong auditory fear memories, is retrieval-dependent, and resistant to spontaneous recovery. Reconsolidation of weak memories is not affected by the expression of GluN2B(E1479Q). (a) Experimental timeline, which includes a second PR-LTM test (spontaneous recovery) 8 days after the initial reactivation. (b) Animals that receive reactivation and infusion with anisomycin displayed significant pharmacologically induced amnesia to the tone 24 hours (*p* < 0.01) and 8 days (*p* < 0.01) following reactivation. (c) Experimental timeline for reconsolidation of weak fear memories. (d) Both the GluN2B(E1479Q) and control animals (GFP) displayed a similar degree of pharmacologically induced amnesia if treated with anisomycin following reactivation. (e) Depiction of the hypothesis that initiation of reconsolidation of strong auditory fear memories can be enabled by increasing GluN2B levels at excitatory synapses. * = *p* < 0.01, *ns* = not significant. Error bars = SEM.

### Increasing synaptic levels of GluN2B in BLA αCaMKII positive neurons using GluN2B(E1479Q) does not influence the reconsolidation of weak auditory fear memories that are capable of being modified via reconsolidation

Next, we sought to determine if increasing GluN2B levels with GluN2B(E1479Q) might further enhance memory destabilization for a weak fear memory, which may lead to an enhanced attenuation of fear when reconsolidation is impaired by post-retrieval intra-BLA anisomycin administration. To accomplish this, we performed a similar experiment to that in Figure 3d, except animals were weak fear conditioned with 3 TSP instead of 10 TSP, (**Figure 4c**). During reactivation, the groups did not significantly differ in their freezing to the tone, [*F*(1,26) = 0.128, *p* = 0.7256], [GFP/Veh *n* = 10, GFP/Aniso *n* = 7, GluN2B(E1479Q)/Veh *n* = 8, GluN2B(E1479Q)/Aniso *n* = 7], (**Figure 4D**). A PR-LTM test was performed 24 hours following reactivation. Two-way ANOVA with repeated measures, [*F*(1,26) = 18.282, *p* = 0.0002], revealed a significant difference between the groups. *Post hoc* analysis revealed that animals receiving anisomycin, regardless of the virus administered, [GluN2B(E1479Q) or GFP], exhibited significantly less freezing compared to saline controls, (*p* < 0.05). Expression of GluN2B(E1479Q) for a weak memory did not lead to an additional enhancement of memory destabilization, indicating that weak fear memories may fully destabilize upon retrieval and/or GluN2B levels are not rate-limiting for the induction of reconsolidation of weak memories.

### Increasing synaptic levels of GluN2B in BLA αCaMKII positive neurons using GluN2B(E1479Q) does not influence the extinction or maintenance of auditory fear memories

GluN2B has been implicated in extinction learning. Blocking the activity of GluN2B-containing NMDARs with ifenprodil within the BLA at the time of extinction has been shown to impair the animal’s ability to extinguish a fear memory (33). However, prior to this study it was unknown whether selectively increasing synaptic GluN2B at the time of extinction learning within BLA αCaMKII positive excitatory neurons would influence extinction learning. To examine this, the BLA of αCaMKII-tTA mice were infused with GFP, GluN2B(WT), or GluN2B(E1479Q) lentiviruses while the animals were on Dox. Animals were fear conditioned with 3 TSP to create weak fear memories. Forty-eight hours later, the animals were taken off Dox for 14 days to enable expression of the lentiviral transgenes. Animals were tested for extinction learning with exposure to 10 tone extinction sessions for three consecutive days (**Figure 5a**). A repeated measures ANOVA for the tone averages of each day did not reveal a difference between groups, [*F*(2,27) = 0.948, *p* = 0.3999], [GFP *n* = 10, GluN2B(E1479Q) *n* = 10, GluN2B(WT) *n* = 10], (**Figure 5b**).

**Figure 5.**
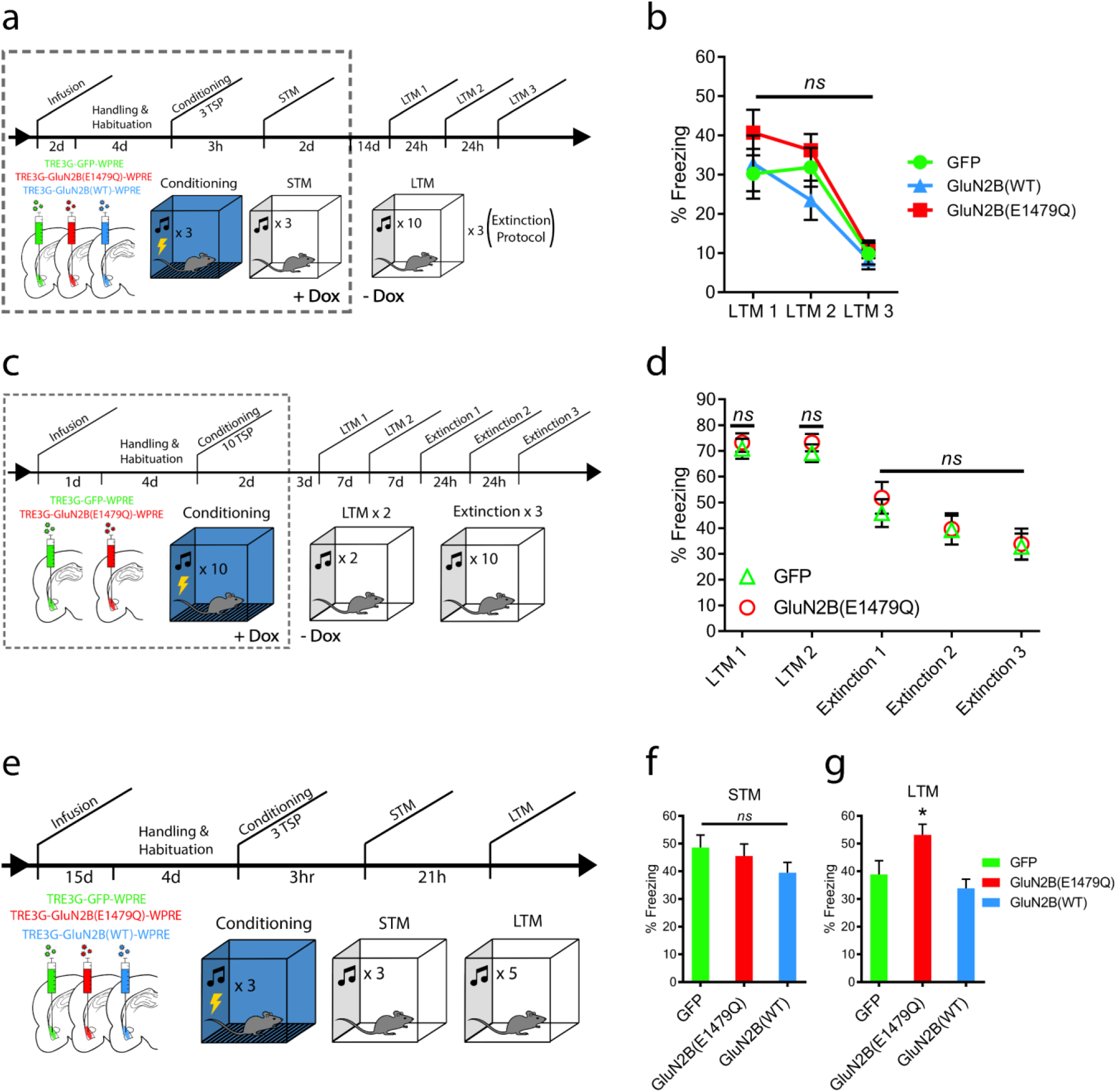
Increasing synaptic GluN2B levels in the BLA utilizing GluN2B(E1479Q) does not affect extinction of strong or weak fear memories, but does enhance consolidation. (a) Experimental timeline for the extinction of weak fear memories. (b) Increasing GluN2B levels in the BLA during extinction training did not enhance or inhibit the ability to extinguish fear. (c) Experimental timeline to test for spontaneous memory loss and extinction of strong fear memories. (d) Expression of GluN2B(E1479Q) within the BLA does not lead to spontaneous loss of a strong fear memory or affect the ability to extinguish fear with extinction training. (e) Experimental timeline to test for the effects of increasing GluN2B levels within the BLA at the time of consolidation. (f) An STM test revealed no difference between GFP, GluN2B(E1479Q), or GluN2B(WT). (g) Animals that received GluN2B(E1479Q) before fear conditioning displayed enhanced consolidation (*p* < 0.01), revealed by an LTM test, while GluN2B(WT) showed similar freezing levels to the GFP control. * = *p* < 0.01, *ns* = not significant. Error bars = SEM.

An additional experiment was conducted to determine if the expression of GluN2B(E1479Q) within BLA neurons interferes with the maintenance and extinction of a strong auditory fear memory over time. Animals were infused with either the GluN2B(E1479Q) or GFP lentiviruses and trained in auditory fear conditioning using 10 TSP while on Dox. All animals were removed from Dox 48 hours following fear conditioning. Five and thirteen days after fear conditioning, LTM tests (2 tones per test) were administered to the animals. Seven days later, the animals were subjected to 3 consecutive days of extinction training (10 tones/day), (**Figure 5c**). There was not a significant difference between the groups during LTM 1, [*F*(1,33) = 0.78, *p* = 0.3828], or LTM 2, [*F*(1,33) = 0.721, *p* = 0.4012], [GFP *n* = 17, GluN2B(E1479Q) *n* = 18]. In addition, no significant differences in freezing between the groups during the three extinction sessions were observed, [*F*(1,33) = 1.017, *p* = 0.3771], (**Figure 5d**).

### Increasing synaptic levels of GluN2B in BLA αCaMKII positive neurons using GluN2B(E1479Q) enhances the consolidation of Pavlovian auditory fear memories

In our final experiment, we examined how the expression of GluN2B(WT) or GluN2B(E1479Q) within BLA excitatory neurons influence the acquisition and consolidation of auditory fear memories. We infused either GFP, GluN2B(WT), or GluN2B(E1479Q) lentiviruses into the BLA of αCaMKII-tTA mice. Fifteen days following infusion, animals were fear conditioned (3 TSP), tested for short-term memory (STM) 3 hours later, and tested for LTM 21 hours after STM (**Figure 5e**). We determined that viral-mediated expression of GluN2B(E1479Q) or GluN2B(WT) prior to and during fear conditioning had no influence on STM, [*F*(2,25) = 1.215, *p* = 0.3125], [GFP *n* = 9, GluN2B(E1479Q) *n* = 9, GluN2B(WT) *n* = 10], (**Figure 5f**), however, the GluN2B(E1479Q) group displayed statistically significantly elevated freezing in response to the tone during the LTM test, [*F*(2,25) = 7.617, *p* = 0.0067], compared to the GFP group, (*p* = 0.0212) and the GluN2B(WT) group, (*p* = 0.002).

## Discussion

Therapeutic methods designed to attenuate maladaptive emotional memories are not currently satisfactorily effective. Disruption of the reconsolidation process has been proposed to be a powerful method to attenuate strong memories in psychopathologies such as PTSD, but numerous studies indicate that strong memories are resistant to retrieval-dependent memory destabilization. Because of this, viable treatments to therapeutically attenuate maladaptive memories by taking advantage of the phenomenon of reconsolidation updating have yet to be fully developed. While others have identified boundary conditions limiting reconsolidation mechanisms, the real challenge is identifying the molecular mechanisms by which these boundaries express themselves in order to develop targeted manipulations to enhance the ability of these circuits to be modified via reconsolidation. Here, we demonstrate that while overexpression of wild-type GluN2B within BLA excitatory neurons is not sufficient to enhance the modification of strong fear memory via reconsolidation mechanisms, the expression of GluN2B(E1479Q) is. These data, for the first time, indicate that increasing *synaptic trafficking* of GluN2B is likely critical for the ability to enhance reconsolidation induction mechanisms – simply increasing mRNA/protein expression may not be sufficient. Therefore, these data refine our understanding of the molecular mechanisms of memory destabilization, and provide evidence for a potential therapeutic target to enhance the induction of reconsolidation in reconsolidation-resistant memories.

Our data indicate that overexpression of GluN2B or GluN2B(E1479Q) does not influence the acquisition of fear memory (i.e., STM), but expression of GluN2B(E1479Q) enhances the consolidation of fear memory (i.e., freezing levels during LTM). Importantly, expression of GluN2B and GluN2B(E1479Q) does not interfere with the expression of fear or sensory perception because STM is intact and freezing levels during reactivation are normal. Notably, ectopic expression of GluN2B or GluN2B(E1479Q) within the BLA does not enhance extinction. It may be that synaptic GluN2B on BLA excitatory neurons is not necessary for, or does not contribute significantly to auditory fear memory extinction. However, given that intra-BLA administration of pharmacological agents that manipulate GluN2B function influence auditory fear memory extinction, it may be that expression of GluN2B on BLA interneurons is necessary for auditory fear memory extinction.

Overexpression of GluN2B(E1479Q), but not GluN2B(WT), enables the retrieval-dependent induction of reconsolidation of a strong fear memory. Our data indicate the likely reason for the failure of overexpression of wild-type GluN2B to enhance the initiation of reconsolidation is that this manipulation simply does not sufficiently increase synaptic GluN2B levels. Other groups have also demonstrated that overexpression of wild-type GluN2B does not significantly increase synaptic GluN2B levels (30, 31). This is in contrast to GluN2A, the overexpression of which does lead to a significant increase in synaptic GluN2A levels (16, 31).

In all, we were able to enhance the modifiability of synaptic connections in a brain region heavily involved in the storage and modulation of fear memories by increasing the synaptic levels of a receptor subunit known to increase synaptic plasticity. Expanding this research to examine agents that increase synaptic trafficking of GluN2B, their involvement in neural circuit stability, and the ability of these manipulations to enhance reconsolidation of modification-resistant memory traces may lead to the development of novel therapeutic strategies to attenuate pathological symptoms and improve the health of patients suffering from a variety of psychiatric disorders.

## Acknowledgements

We would like to thank Saad Akbar for assistance with animal behavior and Manasi Inamdar for assistance with animal behavior and western blot. We would also like to thank Namrata Kumar and Aradhana for their assistance with molecular cloning. This work was supported by the National Institutes of Health (Grants: RMH096202A, RMH109945) and The University of Texas at Dallas.

## REFERENCES CITED

1. Nader K, Schafe GE, Le Doux JE. Fear memories require protein synthesis in the amygdala for reconsolidation after retrieval. Nature. 2000 Aug 17;406(6797):722–6. PubMed PMID: 10963596.

2. Nader K, Schafe GE, LeDoux JE. The labile nature of consolidation theory. Nature reviews Neuroscience. 2000 Dec;1(3):216–9. PubMed PMID: 11257912.

3. Alberini CM. The role of reconsolidation and the dynamic process of long-term memory formation and storage. Frontiers in behavioral neuroscience. 2011;5:12. PubMed PMID: 21436877. Pubmed Central PMCID: 3056265.

4. Tronson NC, Taylor JR. Molecular mechanisms of memory reconsolidation. Nature reviews Neuroscience. 2007 Apr;8(4):262–75. PubMed PMID: 17342174.

5. Przybyslawski J, Sara SJ. Reconsolidation of memory after its reactivation. Behavioural brain research. 1997 Mar;84(1-2):241–6. PubMed PMID: 9079788.

6. McKenzie S, Eichenbaum H. Consolidation and reconsolidation: two lives of memories? Neuron. 2011 Jul 28;71(2):224–33. PubMed PMID: 21791282. Pubmed Central PMCID: 3145971.

7. Tronson NC, Wiseman SL, Olausson P, Taylor JR. Bidirectional behavioral plasticity of memory reconsolidation depends on amygdalar protein kinase A. Nature neuroscience. 2006 Feb;9(2):167–9. PubMed PMID: 16415868.

8. Pitman RK. Will reconsolidation blockade offer a novel treatment for posttraumatic stress disorder? Frontiers in behavioral neuroscience. 2011;5:11. PubMed PMID: 21427793. Pubmed Central PMCID: 3050592.

9. Milton AL, Everitt BJ. The psychological and neurochemical mechanisms of drug memory reconsolidation: implications for the treatment of addiction. The European journal of neuroscience. 2010 Jun;31(12):2308–19. PubMed PMID: 20497475.

10. Suris A, Smith J, Powell C, North CS. Interfering with the reconsolidation of traumatic memory: sirolimus as a novel agent for treating veterans with posttraumatic stress disorder. Annals of clinical psychiatry : official journal of the American Academy of Clinical Psychiatrists. 2013 Feb;25(1):33–40. PubMed PMID: 23376868. Pubmed Central PMCID: 3902858.

11. Taylor JR, Olausson P, Quinn JJ, Torregrossa MM. Targeting extinction and reconsolidation mechanisms to combat the impact of drug cues on addiction. Neuropharmacology. 2009;56 Suppl 1:186–95. PubMed PMID: 18708077. Pubmed Central PMCID: 2635342.

12. Wood NE, Rosasco ML, Suris AM, Spring JD, Marin MF, Lasko NB, et al. Pharmacological blockade of memory reconsolidation in posttraumatic stress disorder: three negative psychophysiological studies. Psychiatry research. 2015 Jan 30;225(1-2):31–9. PubMed PMID: 25441015.

13. Winters BD, Tucci MC, DaCosta-Furtado M. Older and stronger object memories are selectively destabilized by reactivation in the presence of new information. Learning & memory. 2009 Sep;16(9):545–53. PubMed PMID: 19713353.

14. Wang SH, de Oliveira Alvares L, Nader K. Cellular and systems mechanisms of memory strength as a constraint on auditory fear reconsolidation. Nature neuroscience. 2009 Jul;12(7):905–12. PubMed PMID: 19543280.

15. Suzuki A, Josselyn SA, Frankland PW, Masushige S, Silva AJ, Kida S. Memory reconsolidation and extinction have distinct temporal and biochemical signatures. The Journal of neuroscience : the official journal of the Society for Neuroscience. 2004 May 19;24(20):4787–95. PubMed PMID: 15152039.

16. Holehonnur R, Phensy AJ, Kim LJ, Milivojevic M, Vuong D, Daison DK, et al. Increasing the GluN2A/GluN2B Ratio in Neurons of the Mouse Basal and Lateral Amygdala Inhibits the Modification of an Existing Fear Memory Trace. The Journal of neuroscience : the official journal of the Society for Neuroscience. 2016 Sep 7;36(36):9490–504. PubMed PMID: 27605622. Pubmed Central PMCID: 5013194.

17. Ben Mamou C, Gamache K, Nader K. NMDA receptors are critical for unleashing consolidated auditory fear memories. Nat Neurosci. 2006 Oct;9(10):1237–9. PubMed PMID: 16998481. Epub 2006/09/26. eng.

18. Milton AL, Merlo E, Ratano P, Gregory BL, Dumbreck JK, Everitt BJ. Double dissociation of the requirement for GluN2B- and GluN2A-containing NMDA receptors in the destabilization and restabilization of a reconsolidating memory. The Journal of neuroscience : the official journal of the Society for Neuroscience. 2013 Jan 16;33(3):1109–15. PubMed PMID: 23325248. Pubmed Central PMCID: 4241020.

19. Jarome TJ, Werner CT, Kwapis JL, Helmstetter FJ. Activity dependent protein degradation is critical for the formation and stability of fear memory in the amygdala. PLoS One. 2011;6(9):e24349. PubMed PMID: 21961035. Pubmed Central PMCID: 3178530.

20. Hong I, Kim J, Kim J, Lee S, Ko HG, Nader K, et al. AMPA receptor exchange underlies transient memory destabilization on retrieval. Proceedings of the National Academy of Sciences of the United States of America. 2013 May 14;110(20):8218–23. PubMed PMID: 23630279. Pubmed Central PMCID: 3657785.

21. Cull-Candy S, Brickley S, Farrant M. NMDA receptor subunits: diversity, development and disease. Curr Opin Neurobiol. 2001 Jun;11(3):327–35. PubMed PMID: 11399431. Epub 2001/06/12. eng.

22. Prybylowski K, Wenthold RJ. N-Methyl-D-aspartate receptors: subunit assembly and trafficking to the synapse. J Biol Chem. 2004 Mar 12;279(11):9673–6. PubMed PMID: 14742424. Epub 2004/01/27. eng.

23. Sheng M, Cummings J, Roldan LA, Jan YN, Jan LY. Changing subunit composition of heteromeric NMDA receptors during development of rat cortex. Nature. 1994 Mar 10;368(6467):144–7. PubMed PMID: 8139656. Epub 1994/03/10. eng.

24. Monyer H, Sprengel R, Schoepfer R, Herb A, Higuchi M, Lomeli H, et al. Heteromeric NMDA receptors: molecular and functional distinction of subtypes. Science. 1992 May 22;256(5060):1217–21. PubMed PMID: 1350383. Epub 1992/05/22. eng.

25. Lopez de Armentia M, Sah P. Development and subunit composition of synaptic NMDA receptors in the amygdala: NR2B synapses in the adult central amygdala. J Neurosci. 2003 Jul 30;23(17):6876–83. PubMed PMID: 12890782. Epub 2003/08/02. eng.

26. Gardoni F, Caputi A, Cimino M, Pastorino L, Cattabeni F, Di Luca M. Calcium/calmodulin-dependent protein kinase II is associated with NR2A/B subunits of NMDA receptor in postsynaptic densities. J Neurochem. 1998 Oct;71(4):1733–41. PubMed PMID: 9751209. Epub 1998/09/29. eng.

27. Strack S, Colbran RJ. Autophosphorylation-dependent targeting of calcium/ calmodulin-dependent protein kinase II by the NR2B subunit of the N-methyl- D-aspartate receptor. J Biol Chem. 1998 Aug 14;273(33):20689–92. PubMed PMID: 9694809. Epub 1998/08/08. eng.

28. Mayford M, Bach ME, Huang YY, Wang L, Hawkins RD, Kandel ER. Control of memory formation through regulated expression of a CaMKII transgene. Science. 1996 Dec 6;274(5293):1678–83. PubMed PMID: 8939850. Epub 1996/12/06. eng.

29. Holehonnur R, Lella SK, Ho A, Luong JA, Ploski JE. The production of viral vectors designed to express large and difficult to express transgenes within neurons. Molecular brain. 2015;8(1):12. PubMed PMID: 25887710. Pubmed Central PMCID: 4359567.

30. Foster KA, McLaughlin N, Edbauer D, Phillips M, Bolton A, Constantine-Paton M, et al. Distinct roles of NR2A and NR2B cytoplasmic tails in long-term potentiation. J Neurosci. 2010 Feb 17;30(7):2676–85. PubMed PMID: 20164351. Pubmed Central PMCID: 2840640. Epub 2010/02/19. eng.

31. Barria A, Malinow R. Subunit-specific NMDA receptor trafficking to synapses. Neuron. 2002 Jul 18;35(2):345–53. PubMed PMID: 12160751. Epub 2002/08/06. eng.

32. Sanz-Clemente A, Matta JA, Isaac JT, Roche KW. Casein kinase 2 regulates the NR2 subunit composition of synaptic NMDA receptors. Neuron. 2010 Sep 23;67(6):984–96. PubMed PMID: 20869595. Pubmed Central PMCID: 2947143. Epub 2010/09/28. eng.

33. Laurent V, Marchand AR, Westbrook RF. The basolateral amygdala is necessary for learning but not relearning extinction of context conditioned fear. Learning & memory. 2008 May;15(5):304–14. PubMed PMID: 18463174. Pubmed Central PMCID: 2364602.

